# Transcriptomic-based selection of reference genes for quantitative real-time PCR in an insect endosymbiotic model

**DOI:** 10.1101/2023.01.31.526490

**Authors:** Agnès Vallier, Elisa Dell’Aglio, Mariana Galvão Ferrarini, Ophélie Hurtado, Carole Vincent Monégat, Abdelaziz Heddi, Rita Rebollo, Anna Zaidman-Rémy

**Affiliations:** Univ Lyon, INRAE, INSA-Lyon, BF2I, UMR 203, 69621 Villeurbanne, France; Univ Lyon, INSA-Lyon, INRAE, BF2I, UMR 203, 69621 Villeurbanne, France

**Keywords:** real-time PCR, insects, symbiosis, transcriptomics, normalization

## Abstract

Reference genes are a fundamental tool for analyses of gene expression by real-time quantitative PCR (qRT-PCR), in that they ensure the correct comparison between conditions, stages, or treatments. Because of this, selection of appropriate genes to use as references is crucial for proper application of the technique. Nevertheless, efforts to find appropriate, stably expressed transcripts are still lacking, in particular in the field of insect science. Here, we took advantage of a massive transcriptomic high-throughput analysis of various developmental stages of the gut and associated-bacteriomes of the cereal weevil *Sitophilus oryzae* and identified a subset of stably expressed genes with the potential to be used as housekeeping genes from the larva to the adult stage. We employed several normalization techniques to select the most suitable genes among our subset. Our final selection includes three genes - *TAO, YTH3* and *PP12A* - which can also be used to compare transcript abundance at various developmental stages in symbiotic insects, and in insects devoid of endosymbionts (aposymbiotic). Since they are well conserved, these genes have the potential to be useful for many other insect species. This work confirms the interest in using large-scale, unbiased methods for reference gene selection.

## 1 Introduction

Although high-throughput transcriptomic analyses are becoming more affordable and accessible, analysis of a subset of transcripts with quantitative real-time PCR (qRT-PCR) (Higuchi et al., 1992) is still routinely required in laboratory practice, exploratory experiments, experimental validations as well as diagnostics protocols (Kubista et al., 2006). The instrumentation for qRT-PCR is common in molecular biology facilities, along with established guidelines (MIQE guidelines (Bustin et al., 2009)), making it a highly affordable and reliable technique. However, the qRT-PCR accuracy relies on the use of a control method for data normalization that is often achieved by comparing the expression level of target genes against stably expressed transcripts from so-called reference genes, housekeeping genes or constitutively-expressed genes. These transcripts must be stably expressed in the biological samples and experimental conditions that are tested, in order to be used as a comparison to evaluate changes in the expression of the transcripts of interest.

Hence, an appropriate set of qRT-PCR reference genes must be established for each experimental condition, species and tissue examined (Bustin et al., 2009). However, historically, only a small group of highly stably-expressed genes has been used and tested in various conditions and species. A review summarizing recent work on reference genes in 78 insect species from 2008 to 2017 has shown that the majority of reports focused on the same gene candidates, including *Actin, Tubulin, Glyceraldehyde 3-phosphate dehydrogenase (GAPDH*) and the ribosomal gene *18S* (Lü et al., 2018). All these genes displayed great stability and high expression levels in various experimental conditions, tissues and species, but none appeared to be a universal ‘passe-partout’, as highlighted now that transcriptomic analyses are routinely employed in different tissues and at different developmental stages. For example, *GAPDH* (Dveksler et al., 1992) often shows stable expression between insect tissues and treatments but is not often adapted for comparisons between developmental stages (Fu et al., 2013; Pan et al., 2015).

Since the production of transcriptomic data is becoming more and more affordable, it is now possible to choose the best set of reference genes for each species, condition, and treatment, by comparing whole transcriptomes instead of limiting the choice to a set of potential candidates. Examples of this non-aprioristic process are already available for various organisms, including plants (Yim et al., 2015; Liang et al., 2020; Lopes et al., 2021), mammals (Zhang et al., 2020) and human tissues (Caracausi et al., 2017), but not yet for insects, notably, Coleoptera. Around 400 thousand Coleoptera species have been described to date, including terrestrial and aquatic species (Zhang, 2013; Zhang et al., 2018). True weevils, or Curculionidae, are one of the largest animal families and include major agronomic pests, such as the cereal weevil (*Sitophilus* spp.), which is responsible for important damage to fields and stored grains (Longstaff, 1981).

As most insects thriving on a nutritionally unbalanced diet, *S. oryzae* shares a symbiotic relationship with an intracellular Gram-negative bacterium called *Sodalis pierantonius* (Mansour, 1930; Heddi et al., 1999; Oakeson et al., 2014). *S. pierantonius* is housed within specific host cells, the bacteriocytes, forming a unique bacteriome organ attached to the gut during all larval stages (Pierantoni, 1927; Mansour, 1930). Throughout *S. oryzae’s* metamorphosis, the bacteriocytes migrate along the midgut and endosymbionts subsequently infect cells in the apexes of gut caeca (Maire et al., 2020). These coordinated changes result in the formation of multiple bacteriomes along the midgut by the completion of metamorphosis. In these bacteriomes, endosymbionts exponentially increase in number in the emerging adult, before being recycled by an apoptotic-autophagic mechanism following the first week of adult life (Vigneron et al., 2014). Analyzing transcriptomic regulations across weevil’s development is required to fully understand *S. oryzae* and *S. pierantonius* relationship, and to uncover key pathways profitable for integrative pest management strategies. We have recently conducted a thorough analysis of the gut (and associated bacteriomes) transcriptome of *S. oryzae* across its development from larval stages up to bacterial clearance in adults (Figure 1, (Ferrarini et al., 2023)). In this work, we took advantage of the recent transcriptomic profile of *S. oryzae* during development (Ferrarini et al., 2023) to select a set of reliable reference genes for qRT-PCR analyses, with high stability and conservation among insects. The identified transcripts will be of use for the comparison of transcript abundance across multiple developmental stages in *S. oryzae*, gene silencing conditions, and comparisons with artificially-obtained aposymbiotic weevils. These genes also have a great potential for being suitable reference transcripts in other insects, as well as in many other eukaryotes.

**Figure 1:**
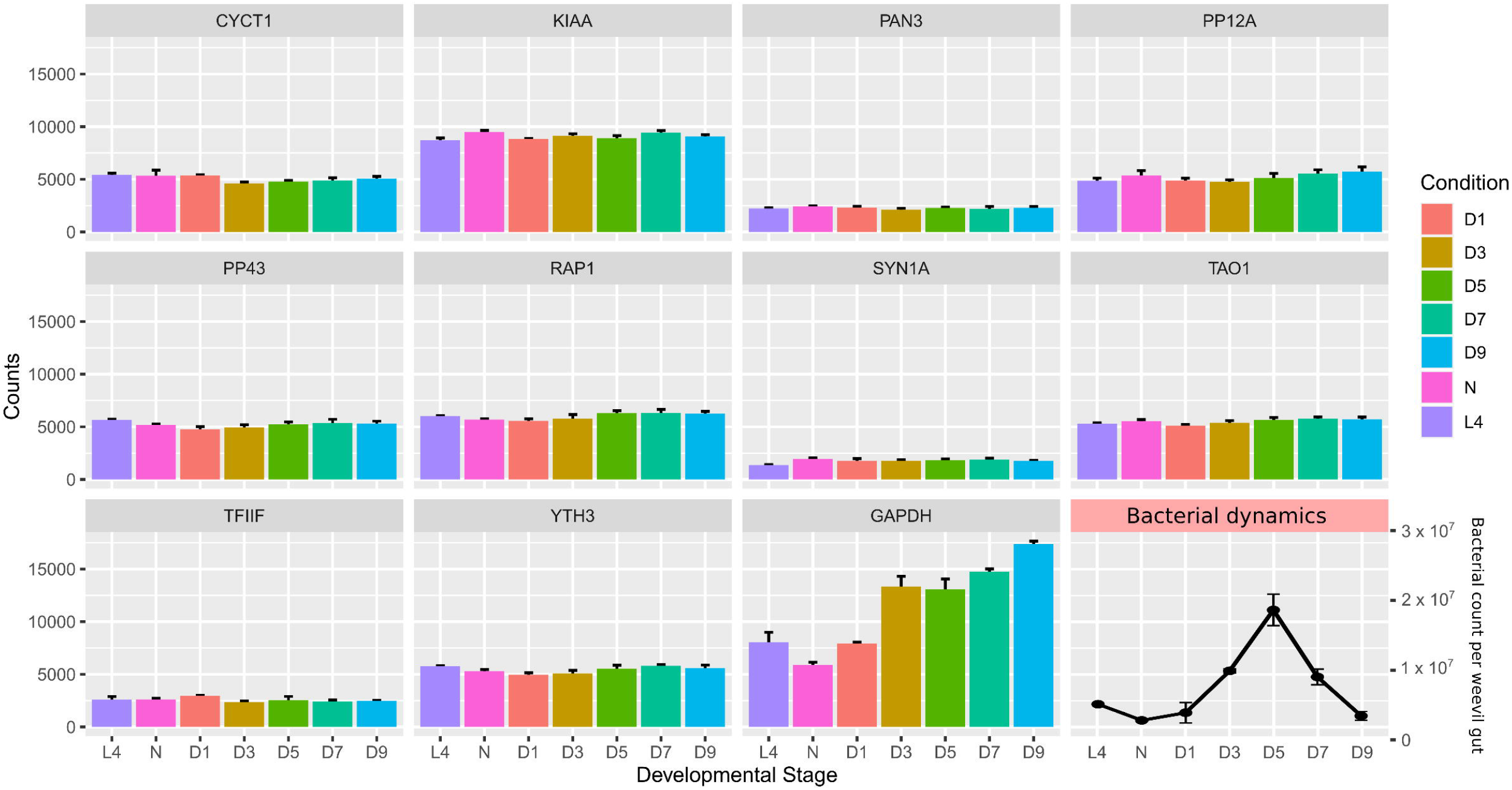
Expression profile of all candidate housekeeping genes in *S. oryzae* gut-bacteriome samples analyzed by transcriptomics at various developmental stages. L4: larval stage 4; P: pupae; D1: 1-day-old adults; D3: 3-day-old adults; D5: 5-day-old adults; D7: 7-day-old adults; D9: 9-day-old adults. Data represent normalized counts at each developmental stage (Ferrarini et al., 2023). Error bars represent standard deviation. Bacterial dynamics is represented in the bottom right corner, adapted from (Dell’Aglio et al., 2022).

## 2 Material and Methods

### 2.1 Insect rearing and sampling

The symbiotic *S. oryzae* population was sampled in Azergues valley, France in 1984, and has been reared in laboratory conditions ever since. The aposymbiotic strain was obtained by heat treatment in 2010, following a protocol described previously (Nardon, 1973a). *S. oryzae* symbiotic and artificially-obtained aposymbiotic weevils were reared in a climate chamber (27.5 °C, 70% relative humidity, no light source) on wheat grains as a food source and substrate for egg laying. In these conditions, the timespan between egg laying and the emergence of adults from the grain is one month for symbiotic weevils, and five weeks for aposymbiotic ones. Adults emerging from the grain are fully formed, although their cuticle is reinforced during the five days following emergence from the grain, together with sexual maturation (Maire et al., 2019). The time between the end of metamorphosis and emergence has been calculated to be around three days (Vigneron et al., 2014). Since our studies are mainly focused on *S. oryzae* transcript levels in the gut, from larval stages to the symbiotic clearance (around day 9 of adulthood), we tested candidate reference genes on gut samples (which contains the bacteriomes) harvested from the last larval stage (L4) to day 9 of adulthood (D9), including pupae (P), and intermediate adult stages (D1, D3, D5, and D7).

For sampling, weevils were immersed in buffer TA (35 mM Tris/HCl, 25 mM KCl, 10 mM MgCl2, 250 mM sucrose, pH 7.5) and dissected under a stereomicroscope. Three biological samples per time point were collected, each composed of five guts and their bacteriomes. All samples were collected on ice in 1.5 mL RNAse-free tubes and stored at −80°C until use.

### 2.2 Transcriptomic analysis

RNAseq datasets were processed in our previous publication (Ferrarini et al., 2023), and raw reads are available at BioProject PRJNA918957. Briefly, quality check was performed with FastQC v0.11.8 (Andrews, 2010), and raw reads were quality trimmed using trim_galore from Cutadapt v0.6.7 (Martin, 2011), then mapped to the *S. oryzae’s* genome (GCA_002938485.2 Soryzae_2.0) using STAR v2.7.3a (Dobin et al., 2013). Then, uniquely mapping reads were counted with featureCounts v2.0.1 (Liao et al., 2014), and mapping quality was assessed by multiqc v1.13 (Ewels et al., 2016). Differential expression analysis was conducted with SarTOOLS, taking advantage of a global linear model from Deseq2 (Love et al., 2014, 2), and log2 fold changes and adjusted p-values were obtained for sequential pairwise comparisons, i.e., L4 vs D1, D1 vs D3 etc, and can be found in the supplementary material of our previous report (Ferrarini et al., 2023). Most graphic outputs are either performed in R, using ggplot2 (Wickham, 2009) or with GraphPad Prism software.

### 2.3 Primer design for candidate reference genes

Primers for each candidate reference gene and *GAPDH* were designed with Primer3 software (Kõressaar et al., 2018, 3), with “best annealing” at 55 °C, amplicon length between 70 and 120 nucleotides (Table 1). Forward and reverse primers were always located in two different exons in order to span at least one intron. In the case of multiple predicted isoforms for each transcript, the primers were designed so that they could amplify all of them. Primer sequences were checked by BLAST (Camacho et al., 2009) to ensure their specificity in the target genome.

**Table 1:**
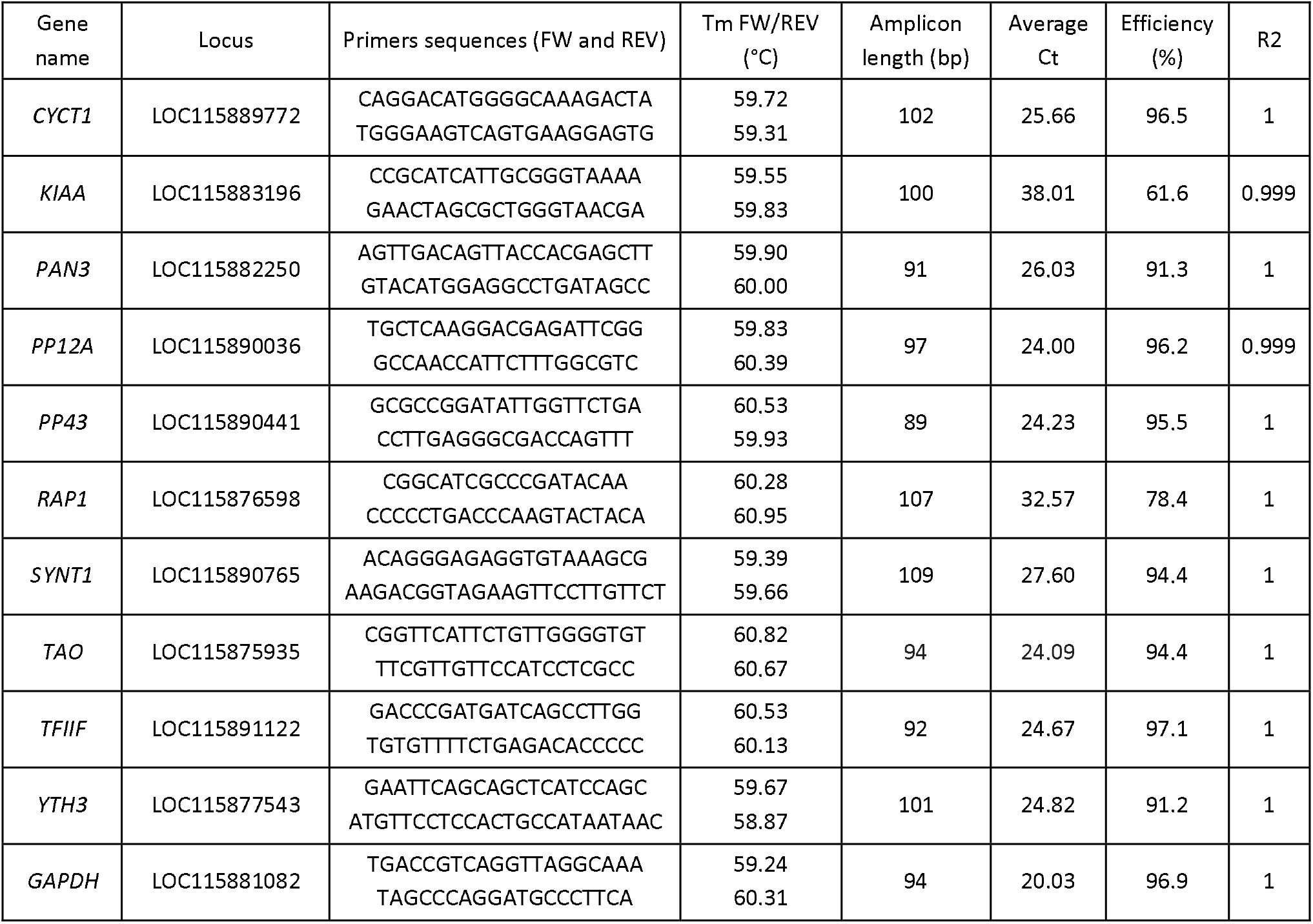
Primers and amplification parameters used for each candidate reference gene selected for qRT-PCR normalization.

### 2.4 RNA extraction and transcript amplification

Total RNA was purified from samples using the RNAqueous Micro kit (Ambion) and its DNAse treatment, following the manufacturer’s instructions. Final RNA concentration was measured with a Nanodrop^®^ spectrophotometer (Thermo Scientific), and RNA quality was checked using agarose gel electrophoresis. Reverse transcription to cDNA was carried out using the iScript™ cDNA Synthesis Kit (Bio-Rad), starting from 1 μg of total RNA. Transcript amplification by qRT-PCR was performed with a CFX Connect Real-Time detection system (Bio-Rad) using the LightCycler Fast Start DNA Master SYBR Green I kit (Roche Diagnostics) as previously described (Maire et al., 2019). Each reaction contained 5 μl of Master Sybr Green mix, 0.5 μl of each primer (10 μM), 1.5 μl RNAse Free water, and 2.5 μl of 1:5 diluted cDNA. Primers for amplification are listed in Table 1. After 5 min at 95 °C, the cycling conditions were: 45 cycles at 95 °C for 10 s, 56 °C for 20 s, and 72 °C for 30 s. A melting curve was obtained at the end of each PCR by heating for 30 s at 66 °C, then increasing the temperature up to 95 °C (increment rates of 0.11 °C/s). For every candidate gene, three biological and two technical replicates were tested for each developmental stage, together with a standard curve made with PCR products as templates, at concentrations spanning from 0.2 fg/μl to 2 pg/μl of amplicon, to calculate the reaction efficiency and ensure linearity of the amplification. The data from each transcript were collected within the same plate.

### 2.5 Candidate reference gene comparisons

To evaluate the reliability of each candidate reference gene, we compared the average cycle threshold (Ct) of each developmental stage and the percentage covariance (CV%) value for each gene (Boda et al., 2009; Sundaram et al., 2019). We then compared the rankings of various web-based algorithms: GeNorm (Vandesompele et al., 2002), the ΔCt method (Silver et al., 2006), BestKeeper (Pfaffl et al., 2004), and Reffinder (Xie et al., 2012). Each candidate reference gene was compared, ranked and evaluated through the tools described.

## 3 Results and discussion

### 3.1 Choice of candidate reference genes based on transcriptome data

To find *S. oryzae* reference genes to be used across gut and bacteriomes development (from larval stage 4 to 9-day-old adults), we scanned a comprehensive transcriptomic dataset spanning *S. oryzae* life cycle (Ferrarini et al., 2023). Candidate reference genes should be stable between conditions, highly expressed, and present transcript isoforms suitable for primer design. There are 17 957 genes in *S. oryzae* (Parisot et al., 2021) and our previous analysis showed that 11 908 were differentially expressed between sequential conditions (Ferrarini et al., 2023). In order to find stable genes across development, we removed genes presenting an adjusted p-value < 0.05 between time points (~1 500 remaining genes), along with a significant log2 fold change lower than −1 or higher than 1 between two consecutive stages, yielding 100 candidate genes. A selection was made by privileging highly-expressed genes (mean expression level higher than 1000), in order to improve qRT-PCR detection, decreasing the candidate list to only 19 genes. Among the resulting pre-selected 19 genes, we visually selected those with the most steady expression along the development. We eliminated genes encoding regulators, which are more likely to change in expression according to the experimental conditions. Among the remaining genes, we selected ones with presumed conserved functions, hence avoiding uncharacterized genes, abundant in the *S. oryzae* genome. These filtering steps resulted in selecting nine potential reference genes for experimental validation (Table 1 and Figure 1). The selected genes spanned various predicted biological functions, to reduce the risk of co-regulation among them and to facilitate the future use of the same genes in other organisms. They included: LOC115875935 – serine/threonine-protein kinase Tao (*TAO*), LOC115876598 – ras-related protein Rap1 (*RAPT*), LOC115877543 – YTH domain-containing family protein 3 (*YTH3*), LOC115882250 – PAN2-PAN3 deadenylation complex subunit Pan3 (*PAN3*), LOC115883196 – transmembrane protein KIAA1109 homolog (*KIAA*), LOC115889772 – cyclin-T1 (*CYCTT*), LOC115890036 – protein phosphatase 1 regulatory subunit 12A (*PP12A*), LOC115890441 – serine/threonine-protein phosphatase 4 regulatory subunit 3 (*PP43*), and LOC115891122 – general transcription factor IIF subunit 1 (*TFIIF*).

Finally, we also included *GAPDH*, a very common reference gene, and the homologue of syntaxin-1 (*SYN1A*), a protein involved in intracellular vesicle traffic, especially in neurons (Zhou et al., 2000), previously selected as a suitable reference gene by (Lord et al., 2010) for *Tribolium castaneum*, the most widely studied member of Coleoptera. Both genes showed little variation between time points and passed all filtering conditions imposed for the other candidate genes (Figure 1). Altogether, a set of 11 candidate genes were then submitted for further analysis.

### 3.2 Expression profiles of candidate reference genes

While the transcriptomic analysis suggests the candidate genes are stable across *S. oryzae’*s gut development, it is important to verify that it is indeed possible to amplify such transcripts in an accurate manner by qRT-PCR. A preliminary PCR test allowed amplification of all predicted transcripts from gut cDNA pools, obtaining fragments compatible with the predicted amplicon sizes. Subsequent qRT-PCR quantification from gut samples (larval stage L4, pupal stage P, and adult stages D1, D3, D5, D7, and D9) was successful for nine out of eleven genes, leading to acceptable efficiency and linearity (Table 1). Two exceptions were the primers for *KIAA* and *RAP1*, which did not lead to acceptable fragment amplification under standard conditions (efficiency values of 61.6 and 78.4, respectively; average Ct values of 38.01 and 32.58, respectively); and were therefore excluded from the list of potential candidates (Table 1). Moreover, although successful, *YTH3* expression levels appear lower than expected based on the transcriptomic analysis. Discrepancies between amplification in qRT-PCR and normalized read counts could be due to low primer efficiency, a tendency to primer dimerization, or suboptimal amplification of transcripts in our standardized qRT-PCR protocol, as well as differences in normalization strategies between the two techniques. One should also note that for qRT-PCR, larval samples consisted of bacteriomes and whole guts, while in the previous transcriptomic dataset on which the candidate reference gene selection was based, only larval bacteriomes were dissected (Ferrarini et al., 2023).

The mean Ct values of amplicons for each candidate reference gene at each developmental stage are reported in Figure 2. The gene with the lowest average Ct was *GAPDH* (mean for all stages: 20.0), followed by *PP12A* (mean for all stages: 23.99) and *PP43* (mean for all stages: 24.66). All average Ct values are considered low (*GAPDH*) or medium (all other genes) and are therefore suitable for estimating gene expression of various target genes with different expression profiles. Statistical analysis of Ct values revealed that the mean expression level for the majority of the genes is subjected to mild variations between developmental stages (see asterisks in Figure 2), except for *GAPDH* and *SYN1A*.

**Figure 2:**
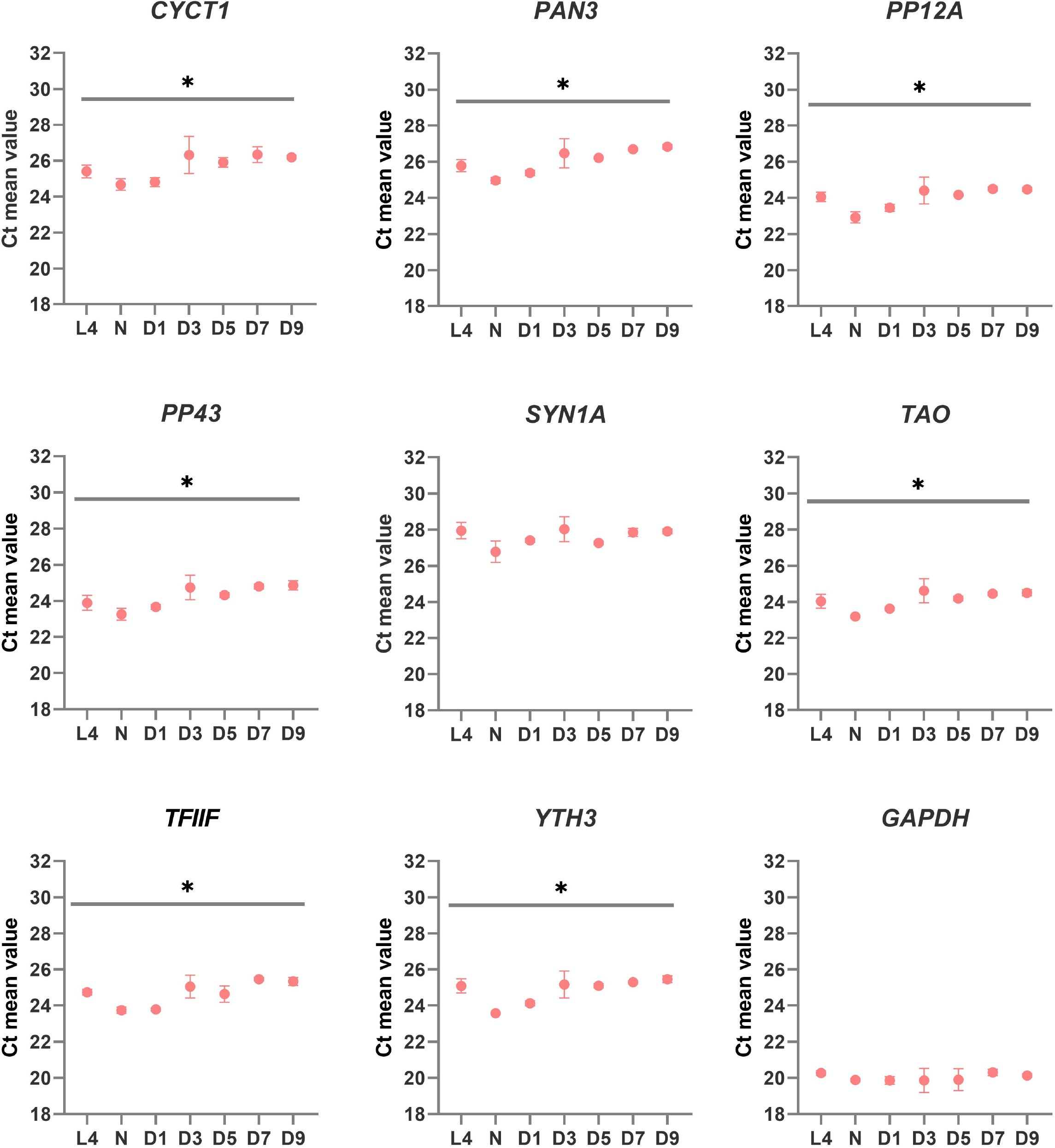
qRT-PCR expression profile of candidate reference genes in *S. oryzae* gut-bacteriome samples. Graphs represent the mean Ct value for each developmental stage. One-way ANOVA was performed to assess differences between the means of all developmental stages. Asterisks denote statistical significance (ANOVA with Kruskal-Wallis test, * = p ≤ 0.05). Error bars represent standard deviation.

### 3.3 Evaluation of expression stability and ranking of candidate reference genes

To identify the most suitable reference gene to compare transcript expression across *S. oryzae* development, we relied on the comparison of statistical methods recently conducted by Sundaram and colleagues (Sundaram et al., 2019). After describing the merits and limitations of various comparison algorithms, the authors suggested the following as the best pipeline for the identification of reliable reference genes: i) visual estimation of expression variation across stages, ii) analysis of CV% values ((Boda et al., 2009); a cut-off of 50 is recommended); then iii) ranking of the remaining candidate reference genes by the NormFinder algorithm (Andersen et al., 2004).

Since the visual estimation of expression variation across stages was already performed and showed mild variation for most genes (Figure 2), we calculated the CV% value for each candidate reference gene. The CV% values of four out of nine genes (*YTH3, CYCT1, PAN3* and *TFIIF*) across all developmental stages are indeed higher than 50 (Figure 3A), reflecting variations observed in particular at the pupal stage (Figure 2). When only adult stages were considered (Figure 3B), only two genes still displayed CV% values higher than 50 (*CYCT1* and *TFIIF*). After excluding genes with CV% higher than 50, we ran the NormFinder algorithm together with other common reference gene ranking algorithms: GeNorm (Vandesompele et al., 2002), the ΔCt method (Silver et al., 2006), BestKeeper (Pfaffl et al., 2004), and Reffinder (Xie et al., 2012). Rankings according to all prediction tools are reported in Table 2 for all stages, and in Table 3 for adult stages only. Overall, each of these methods provided good stability values for all the preselected candidate genes, with a consensus for *TAO* and *SYN1A* at all stages and for *TAO, PP12A* and *YTH3* in adult stages only.

**Figure 3:**
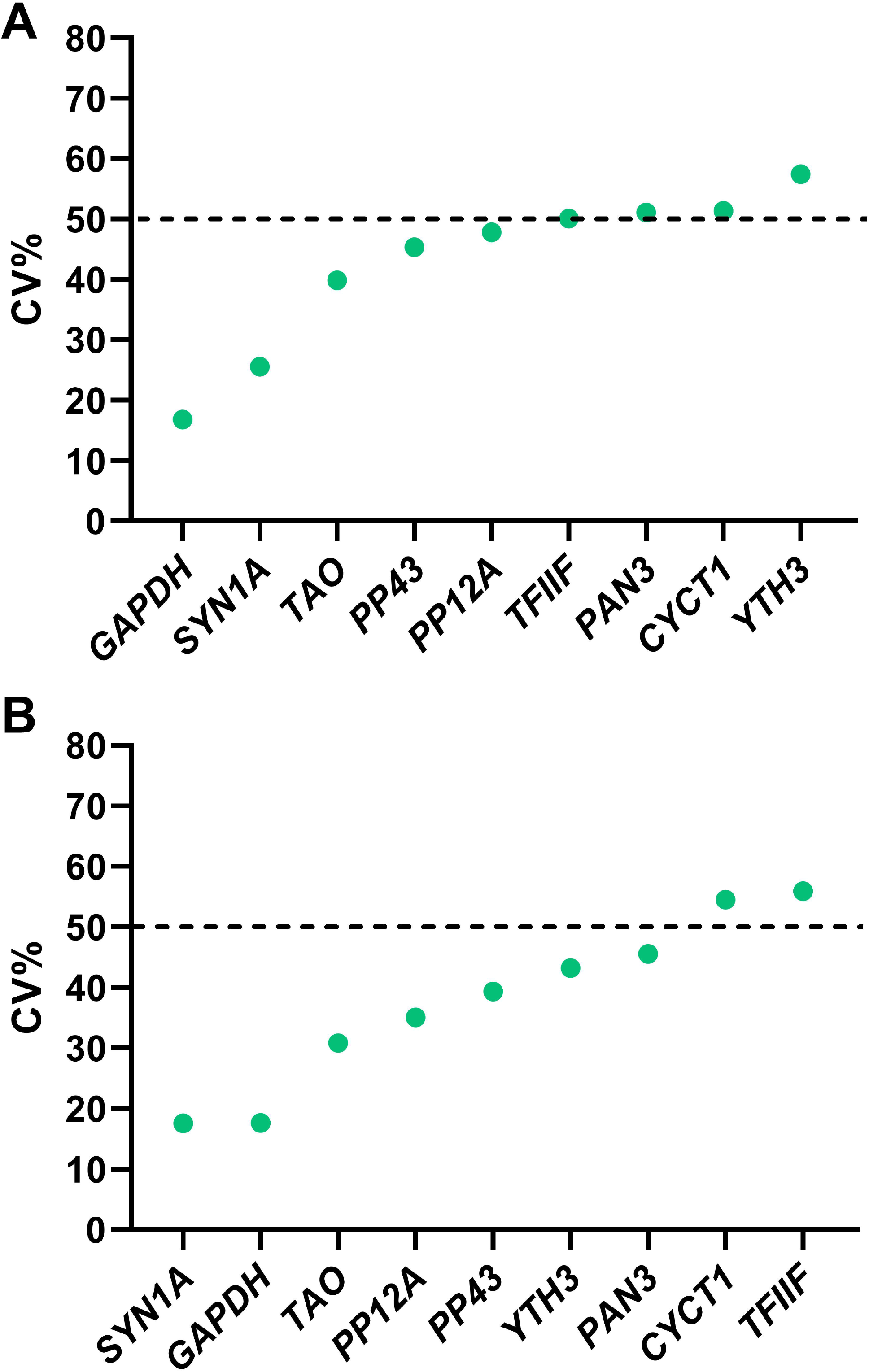
CV% values for the selected candidate reference genes, calculated on the linearized 2-Ct converted Ct values at all *S. oryzae* developmental stages considered in Figure 2. A) CV% measured considering all developmental stages. B) CV% calculated considering only adult stages. Only genes with a CV% lower than 50 were selected for further analysis.

**Table 2:**
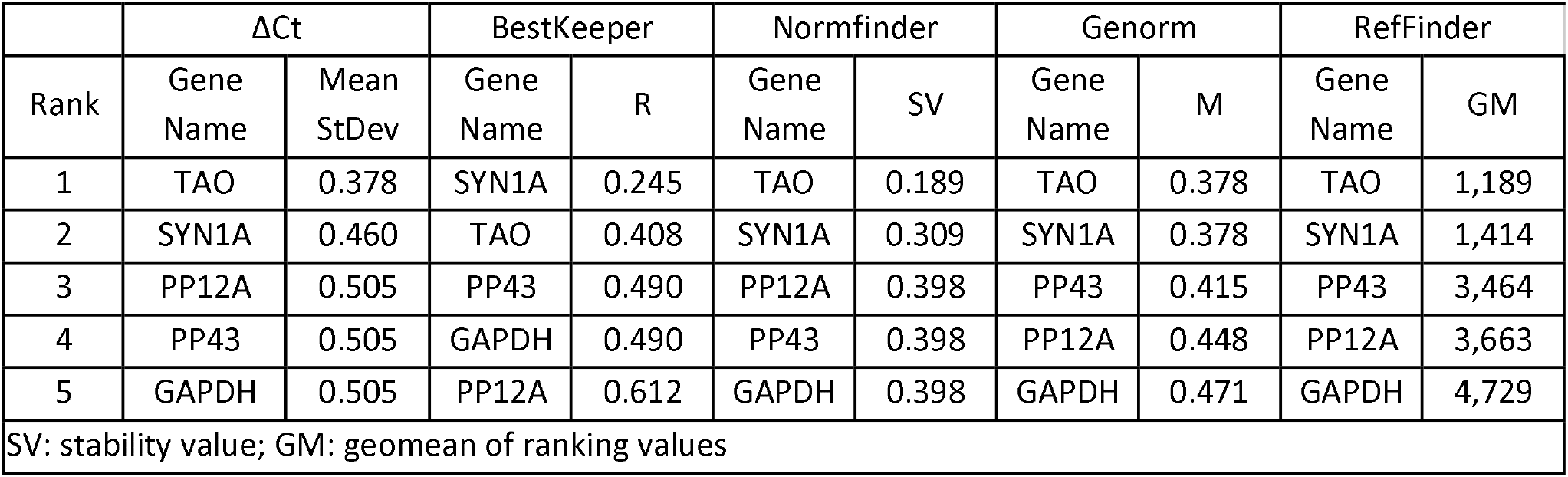
Ranking of candidate reference genes for various *S. oryzae* developmental stages, according to the ΔCt method, BestKeeper, NormFinder, GeNorm, and Reffinder. The rankings were obtained for all developmental stages.

**Table 3:**
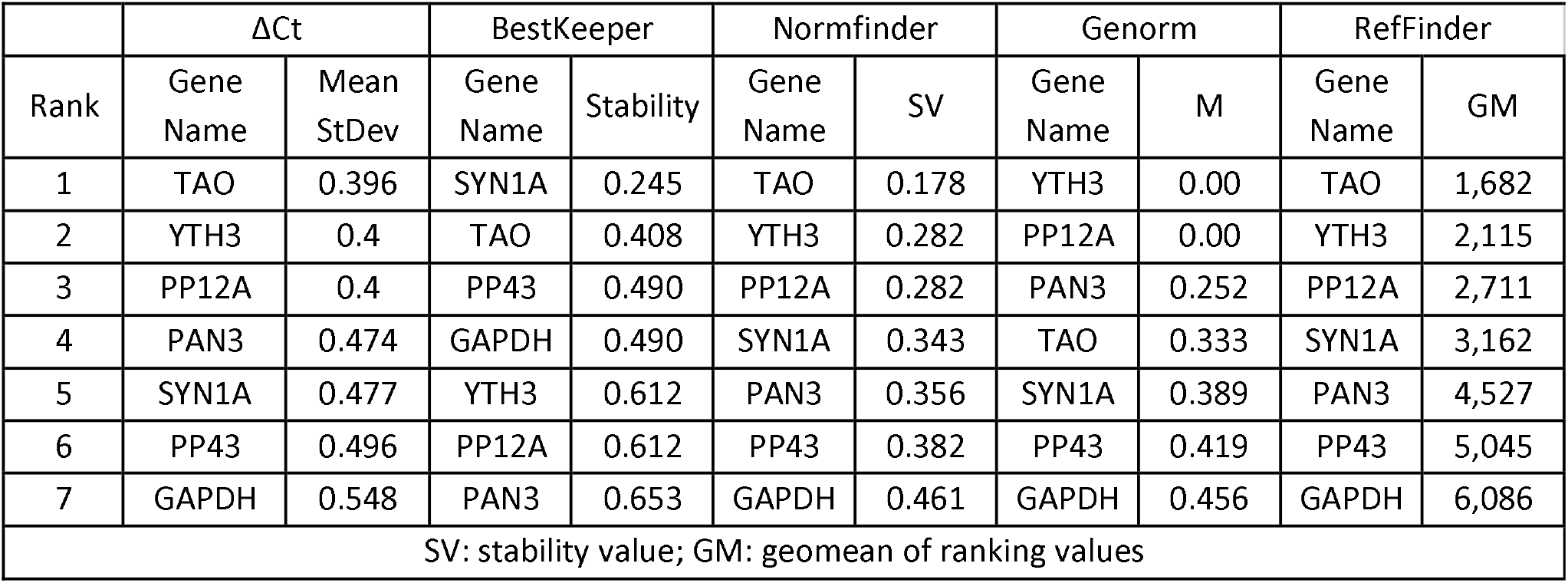
Ranking of candidate reference genes for various *S. oryzae* developmental stages, according to the ΔCt method, BestKeeper, NormFinder, GeNorm, and Reffinder. The rankings were obtained for adult stages only.

### 3.4 Determination of the minimum number of reference genes for qPCR normalization

The pairwise variation value (V) between the ranked genes was calculated with the geNorm algorithm and used to estimate the stability of the normalization factor with the addition of one normalization gene at a time. The cut-off value for pairwise variation used was 0.15 (Vandesompele et al., 2002), meaning that as soon as the value of V drops below 0.15 the number of reference genes was considered suitable for correct standardization of gene expression. The V value for the set of candidate reference genes was always lower than 0.15 (Figure 4). This indicates that two reference genes, which is the minimal number of reference genes accepted by the MIQE guidelines (Bustin et al., 2009), are sufficient for standardization of gene expression across *S. oryzae* development in the gut, from the L4 larval stages to D9 adults. It also shows that any combination of the proposed nine reference genes is suitable for normalization.

**Figure 4:**
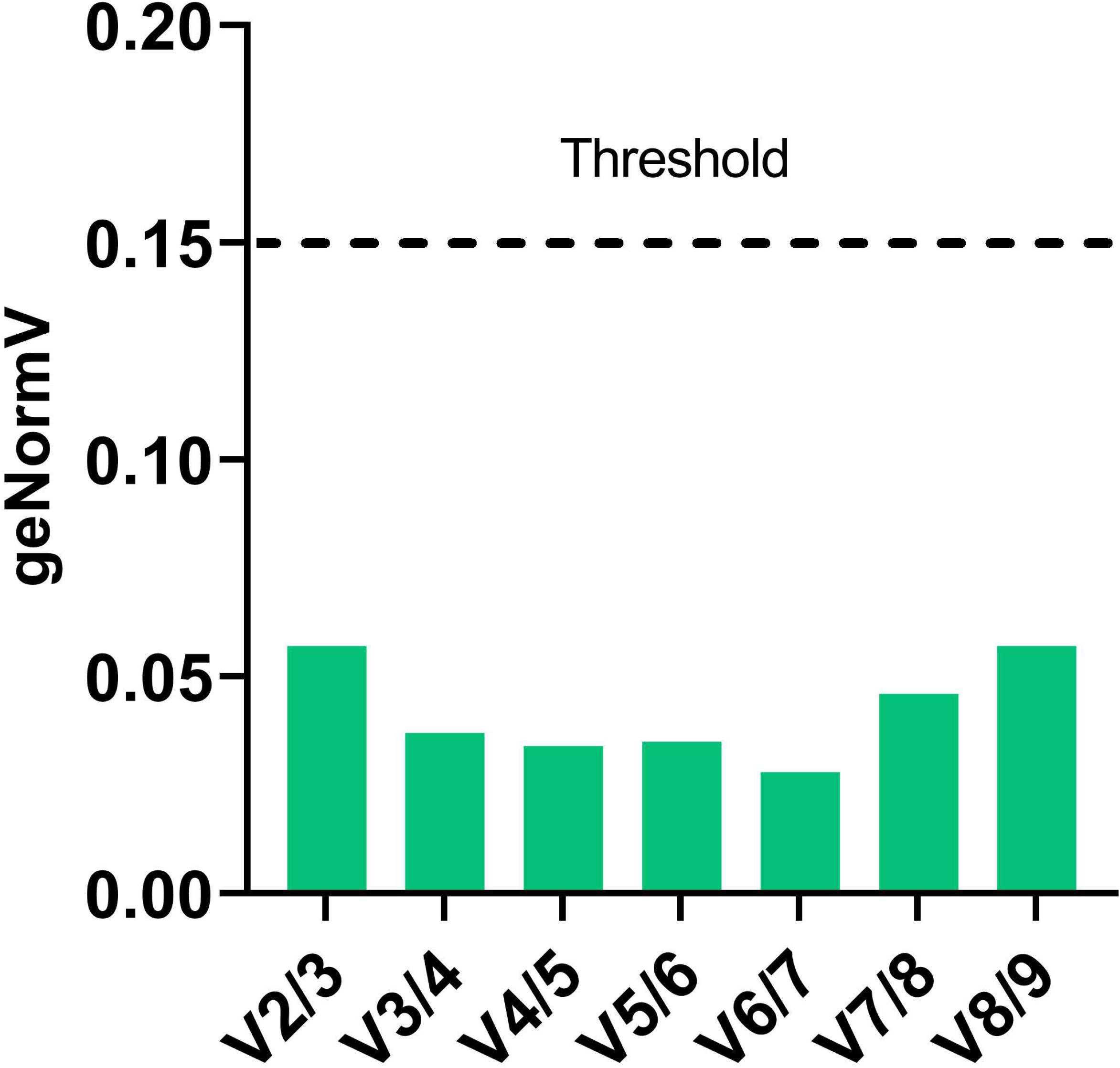
Optimal number of reference genes for normalization in *S. oryzae* at various developmental stages. The pairwise variation values obtained from geNorm analyses were used to determine the minimum number of reference genes for normalization. The value of Vn/Vn+1<0.15 indicates that the minimum number of reference genes to be used for qPCR data normalization is n.

### 3.5 Assessment of candidate reference genes for transcript comparisons between symbiotic and aposymbiotic insects

Weevils artificially deprived of symbionts by heat treatment (aposymbiotic) do survive under laboratory standard conditions, although their fitness is partially compromised, e.g. the larval development takes longer, fertility is lower (Nardon, 1973b), and aposymbiotic adults are unable to fly (Nardon et al., 1998). In aposymbiotic weevils, the insect cuticle, which is derived from aromatic amino acids mainly supplied by the bacteria, does not fully develop but remains thinner and more delicate (Vigneron et al., 2014). In order to pinpoint symbiosis-specific transcriptional changes, the transcriptomic profiles obtained for symbiotic weevils (Ferrarini et al., 2023) should be compared with the expression profiles obtained at the same stage in aposymbiotic weevils. We took advantage of previously transcriptomic datasets produced from adult symbiotic and aposymbiotic weevils at different adult stages, to compare the expression of all candidate genes between symbiotic stages (Figure 5, (Ferrarini et al., 2023)). Differential expression analysis of all nine candidate genes between symbiotic and aposymbiotic weevils, shows only three genes are equally expressed between symbiotic conditions: *PAN3, TAO* and *YTH3*. Two of these genes were chosen as suitable candidates for symbiotic analysis, *TAO* and *YTH3* and should constitute a good set for symbiotic/aposymbiotic comparisons.

**Figure 5:**
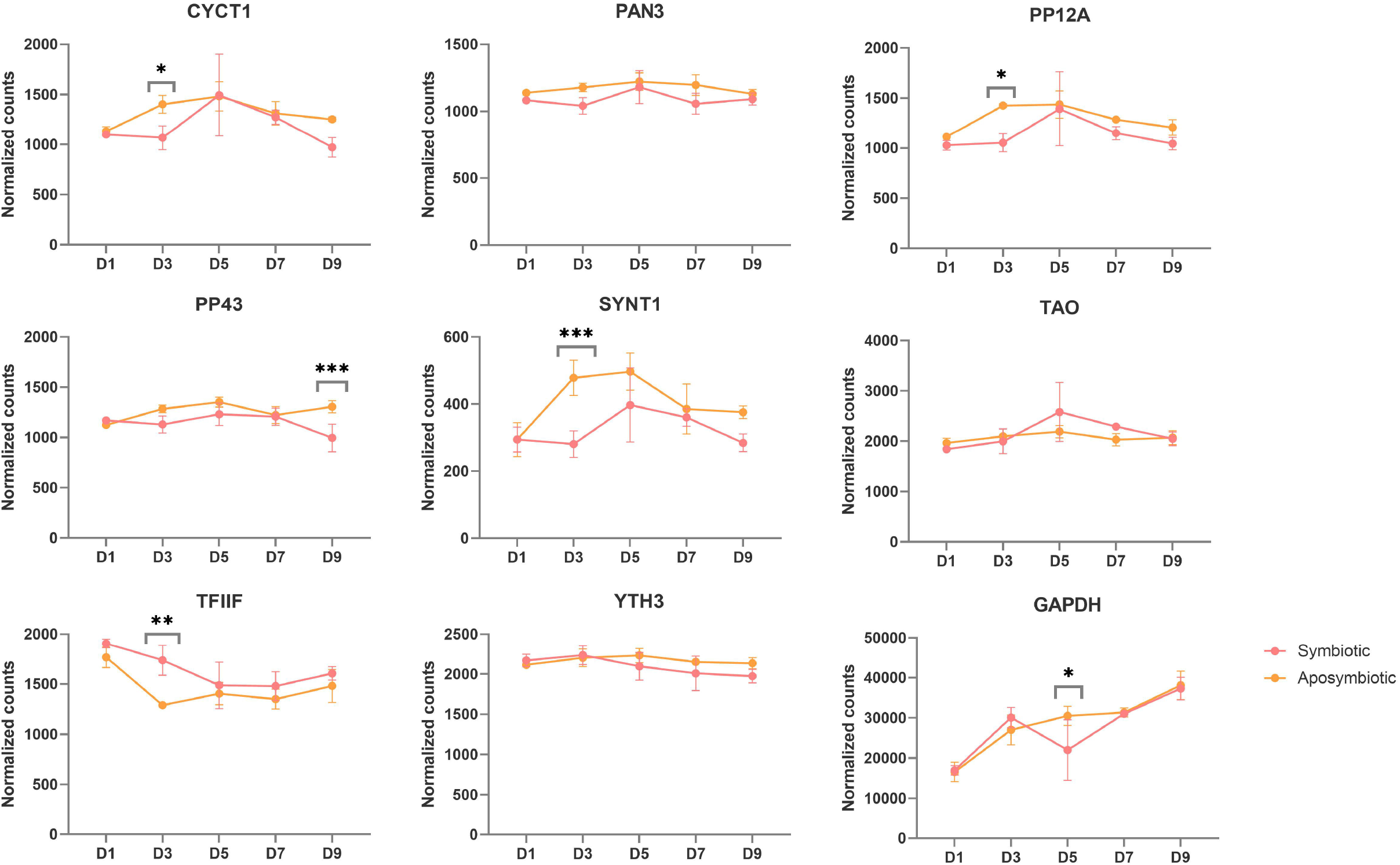
Normalized counts of selected candidate reference genes from previous RNAseq data obtained from symbiotic and aposymbiotic weevils (Ferrarini et al., 2023). Stars denote the adjusted p-value between symbiotic and aposymbiotic weevils at a given time point, upon DESEQ2 analysis (see methods). * adjpvalue < 0.05, ** <0.005, *** < 0.0005, **** < 0.00005. Three genes show no differences between symbiotic and aposymbiotic weevils at any given time point and constitute good candidates for symbiotic/aposymbiotic qRT-PCR comparisons: *pan3, tao* and *yth3*.

## 4 Conclusion

In this work, we relied on transcriptomic data to select a set of nine new candidate reference genes for *S. oryzae*, which we compared with *GAPDH*, a very common reference gene (Bustin, 2000), and *SYN1A*, which has been previously used as a reference gene for *T. castaneum* (Lord et al., 2010). By combining differential expression data filtering, visual inspection, CV% analysis and a variety of gene stability ranking algorithms, we were able to select stable gut transcripts in *S. oryzae* adult stages, but also for stages spanning from larvae to adult, including metamorphosis. In particular, all the preselected candidate genes scored well in all ranking systems, with a general preference for *TAO* and *SYN1A* when all developmental stages (from larval stage 4 to 9-day-old adults) were considered, and for *TAO, PP12A* and *YTH3* in adult stages only. Interestingly, for all the various algorithms used to rank the reliability of reference genes, the newly-selected genes were always more suitable than the highly popular *GAPDH*, although the latter ranked better in terms of CV% values.

Our selection approach was guided by the stability of expression levels of various transcripts and their abundance in transcriptomic data (Ferrarini et al., 2023), and on the presumed conserved, housekeeping function of each gene on the basis of the available literature and sequence alignments with homologs from other species. Indeed, all the candidates that were successfully quantified by qPCR, and in particular *TAO* and *PP12A*, which were chosen for comparing symbiotic and aposymbiotic weevils, have been previously associated with important cell functions, such as DNA repair and RNA transcription, suggesting they are likely good candidates as reference genes for many other tissues and organisms. In mammals, the homologue of TAO (Taok1 in *Rattus norvegicus*) is involved in the G-protein signalling cascade as a serine/threonine-protein kinase active on MAP2K3, MAP2K6 and MARK2. It regulates DNA damage response, apoptosis and cytoskeleton stability (Hutchison et al., 1998; Timm et al., 2006). As these processes are well-conserved also in insects, it is quite likely that *S. oryzae TAO* protein fulfills the same function as the mammalian homolog. On the other hand, PP12A (for Protein Phosphatase 1 Regulatory Subunit 12A, or *PPP1R12A* in *Homo sapiens*), is a subunit of the myosin phosphatase complex responsible for the interaction between actin and myosin (Takahashi et al., 1997; Hughes et al., 2020). The Coleoptera sequence only partially aligns with the *Drosophila* and mammalian sequences (in correspondence to the phosphatase domain located at the N-terminus), suggesting evolutionary divergence and hence a potentially unknown function.

The other genes investigated include: a Ser/Thr phosphatase (PP43, or Protein Phosphatase 4 subunit 3), first identified in yeast as the regulatory component of a Ser/Thr phosphatase and conserved in mammals and *Drosophila* (Gingras et al., 2005; Karman et al., 2020), along with plants (Kataya et al., 2017); an animal-specific transcription initiation factor (TFIIF), part of the pre-initiation complex (Aso et al., 1992; Luse, 2012); the transcriptional regulator PAN3 (Poly-A Nuclease protein 3), which is found in all eukaryotes and is part of a poly-A degradation complex (Brown et al., 1996; Garneau et al., 2007); another animal transcriptional regulator YTH3 (YT521-B Homology 3), involved in the degradation of N6-methyladenosine (m6A)-containing mRNA and non-coding RNAs (Zaccara and Jaffrey, 2020); and finally the *S. oryzae* CYCT1 protein (for Cyclin T1), composed of a cyclin conserved domain at the N terminus, conserved among all animals (Shim et al., 2002), and a long C-terminal with unknown function only conserved among Coleoptera.

In conclusion, we identified stable reference genes across *S. oryzae* development with potential application in various other animal species. This work also demonstrates the potential of transcriptomic data as a guide for reference gene selection to obtain comparable or better performances in comparison to traditional ones - a strategy still neglected in insect studies.

## 6 Author Contributions

AZR and RR conceived the study with the help from AH; MGF performed the gene selection from high throughput transcriptomic data, with inputs from CVM, AV, EDA, AH and AZR; AV, EDA, and OH produced and treated the gene expression data; AV, EDA and RR performed the statistical analysis; AZR, RR, AV, ED, AH and EDA wrote the article. All authors read and approved the manuscript.

## 8 Data Availability Statement

The datasets analyzed for this study can be found at the following BioProject PRJNA918957 (http://www.ncbi.nlm.nih.gov/bioproject/918957).

